# A Functional Menadione Biosynthesis Pathway is Required for Capsule Production by *Staphylococcus aureus*

**DOI:** 10.1101/2021.06.02.446699

**Authors:** Dina Altwiley, Tarcisio Brignoli, Andrew Edwards, Mario Recker, Jean Lee, Ruth C. Massey

## Abstract

*Staphylococcus aureus* is a major human pathogen that utilises a wide array of pathogenic and immune evasion strategies to cause disease. One immune evasion strategy, common to many bacterial pathogens, is the ability of *S. aureus* to produce a capsule that protects the bacteria from several aspects of the human immune system. To identify novel regulators of capsule production by *S. aureus* we applied a genome wide association study (GWAS) to a collection of 300 bacteraemia isolates that represent the two major MRSA clones in UK and Irish hospitals: CC22 and CC30. One of the loci associated with capsule production, the *menD* gene, encodes an enzyme critical to the biosynthesis of menadione. Mutations in this gene that result in menadione auxotrophy induce the slow growing small-colony variant (SCV) form of *S. aureus* often associated with chronic infections due to their increased resistance to antibiotics and ability to survive inside phagocytes. Utilising such an SCV we functionally verified this association between *menD* and capsule production. Although the clinical isolates with polymorphisms in the *menD* gene in our collections had no apparent growth defects, they were more resistant to gentamicin when compared to those with the wild-type *menD* gene. Our work suggests that menadione plays a critical role in the production of the *S. aureus* capsule, and that amongst clinical isolates polymorphisms exist in the *menD* gene that confer the characteristic increased gentamicin resistance, but not the major growth defect associated with SCV phenotype.

## Introduction

As a successful human pathogen *Staphylococcus aureus* has evolved many mechanisms to evade host immunity, including the production of a polysaccharide capsule that protects the bacteria from killing by phagocytes [1-3]. The enzymes responsible for the biosynthesis of this capsule are encoded within a multi-gene locus (*cap*) that has both highly conserved and variable genes responsible for a range of capsule serotypes [2, 4]. The importance of capsule production to the ability of *S. aureus* to cause disease has been demonstrated in many animal models, and as a result it was a target of an anti-staphylococcal vaccine attempt, albeit unsuccessful in clinical trials in humans [5]. Recent population-based analysis of human isolates may partially explain the lack of success of this vaccine in clinical trials, as it found significant variability in the amount of capsule produced by clinical isolates [6, 7]. Although there are associations between the levels of capsule production and increased patient mortality [8], that capsule negative variants are frequently isolated from patients suggests that capsule production is not critical for survival in humans or the ability to cause disease.

Using traditional molecular approaches, several regulators of the expression of the *cap* locus have been identified such as Agr and MgrA [9, 10]. However, the existence of variability across a collection of isolates can facilitate alternative approaches to the identification of novel regulators through the use of genome wide association (GWAS) approaches [8, 11-14]. These have the added benefit of allowing a greater understanding of the role and relevance of these regulators in the natural environment of the human host [8, 11]. A previous application of this approach to a collection of community-acquired methicillin resistant *S. aureus* (MRSA) USA300 isolates identified several conserved mutations within the *cap* locus as responsible for variability in capsule production [6]. Here, we sought to extend this approach to a collection of healthcare-acquired MRSA representing the two major clones circulating in UK and Irish hospitals, clonal complexes 22 and 30 (CC22, CC30).

We observed a high level of variability with regards to capsule production within our collection of clinical isolates. Interestingly, no polymorphisms within the *cap* locus were identified as associated with this phenotype, although several loci distal to the *cap* locus were associated with capsule production. One of these genes, *menD*, encodes an enzyme critical to the biosynthesis of menadione [15]. Mutations in this gene have been shown in many studies to be responsible for an alternative means utilized by *S. aureus* to both resist the effect of antibiotics and evade clearance by phagocytes by switching to the slow growing small colony variant (SCV) or persistor phenotype [15-17]. The expression of many virulence factors is reduced when the bacteria switch to SCVs, including the production of cytolytic toxins [18]; however, there are contradictory reports on what effect this switch has on capsule production [19-21]. In this study we explore the link between the SCV phenotype and capsule production and conclude that the link is dependent upon the specific pathway that becomes mutated during the switch to the SCV form.

## Methods

### Bacterial strains and culture conditions

Bacterial strains used are listed in Table 1. Bacterial strains were routinely stored at -80°C in 15% glycerol/broth stocks until required. Unless stated otherwise, *S. aureus* strains were streaked onto Tryptic Soy agar (TSA) and single colonies transferred to 5 mL Tryptic Soy broth (TSB). All bacterial cultures were propagated in a shaking incubator for 18 h at 37°C at 180 rpm.

### *In vitro* capsule production quantification

Colonies of the *S. aureus* strains were grown overnight in Tryptic Soy Agar (TSA) at 37°C and these were transferred to nitrocellulose (NC) membranes by laying the membrane on the surface of each plate. After two min, the membranes were placed bacteria side up in a clean petri dish and baked for 15 min at 60°C. To remove excess bacteria from the filters, membranes were washed three times in PBS and the proteins removed by incubating the filters in trypsin solution for 1 h at 37°C. Membranes were then rinsed and blocked in Bovine Serum Albumin (BSA) for 1 h, and washed 3X in PBS with 0.05%Tween. The membranes were incubated for 1 h in diluted anti-cap antiserum 1:1000 - 1:3000 (5-15 μL:15mL PBS) at room temperature with gentle agitation. Filters were washed 3X for 3 min each with PBS/Tween. Protein G-HRP conjugate was diluted in PBS/Tween to a 1:5000 dilution and incubated for 1 h at room temperature with gentle agitation. The membranes were washed 3X for 3min each with PBS. Finally, the reactivity of the colonies was detected using the Opti-4CN Substrate Kit (BIORAD), according to manufacturer instructions. The clinical isolates were scored visually by three individuals as 0, 1 or 2, where 0 indicated no capsule detection, 1 a medium level of capsule detection and 2 a high level of capsule detection.

### GWAS

Genome-wide association mapping was conducted using a generalized linear model, with capsule production as the quantitative response variable. We accounted for bacterial population substructure by adding to the regression model the first two component from a *principal component decomposition* of SNP data for each set of clinical samples (CC22 and CC30). The first two components accounted for 32% and 40% of the total variance for CC22 and CC30, respectively. In both cases, three distinct clusters were identified. We further considered a third model where we used cluster membership as covariates in our regression model, where clusters were defined using K-means clustering analysis (setting K=3); this, however, yielded identical results to the one based on PCA components. In total, 2066 (CC22) and 3189 (CC30) unique SNPs were analyzed, the majority of which were subsequently filtered out for exhibiting a minor allele frequency (maf) of <0.03, reducing the data to 378 and 1124 SNPs, respectively. Reported P-values are not corrected for multiple comparison; Sidak corrected significance thresholds are indicated in the Manhattan plots.

### mRNA extraction

The bacteria were grown on TSB at 37°C incubated in shaking incubator for 18h. After the incubation, RNA was extracted by Quick-RNA Fungal/Bacterial Miniprep Kit (Zymo Research) according to the manufacturer’s instructions. RNA integrity was checked by running 5 μL aliquot of the RNA on a 1% agarose gel and observing the intensity of the ribosomal RNA (rRNA). RNA samples were treated by TURBO™ DNase (Invitrogen) to eliminate any genomic DNA contamination. To verify that the samples were free from any DNA contamination, RNA samples were subjected to RT-qPCR alongside with a no template control (NTC) and 2.5 ng of a known genomic DNA and threshold rate were compared.

### Quantitative reverse transcriptase (RT-qPCR)

To quantify the expression of the *capE* gene of the wild type and the mutants, RT-qPCR was performed using *gyrB* as a reference gene. Complementary DNA (cDNA) was generated from mRNA using qScript® cDNA Synthesis Kit and following the manufacture’s protocol (Quantabio) and the cDNA was used as a template for the qPCR reaction. Primers used are listed in table 2. The reverse-transcriptase PCR (RT-PCR) was performed as follows: 10 μl 2x SensiFAST SYBR Mix, 0.8 μl of 10 μM forward primer, 0.8 μl of 10 μM μl reverse primer, 1 μl cDNA and RNase-free water up to a total of 20 μl volume. The PCR condition cycles consisted of initial denaturation at 95°C for 2 min followed by 40 cycles of denaturation at 95°C for 10 s, annealing at 55°C for 60 s and extension at 72°C for 10 s. RT-PCR was carried out in triplicate for each sample and 3 biological repeats or more using the primers listed in Table 1. The ratio of *capE* and *gyrB* transcript number was calculated using the using the 2^-(ΔCt ply – ΔCt recA)^ method.

**Table 2:**
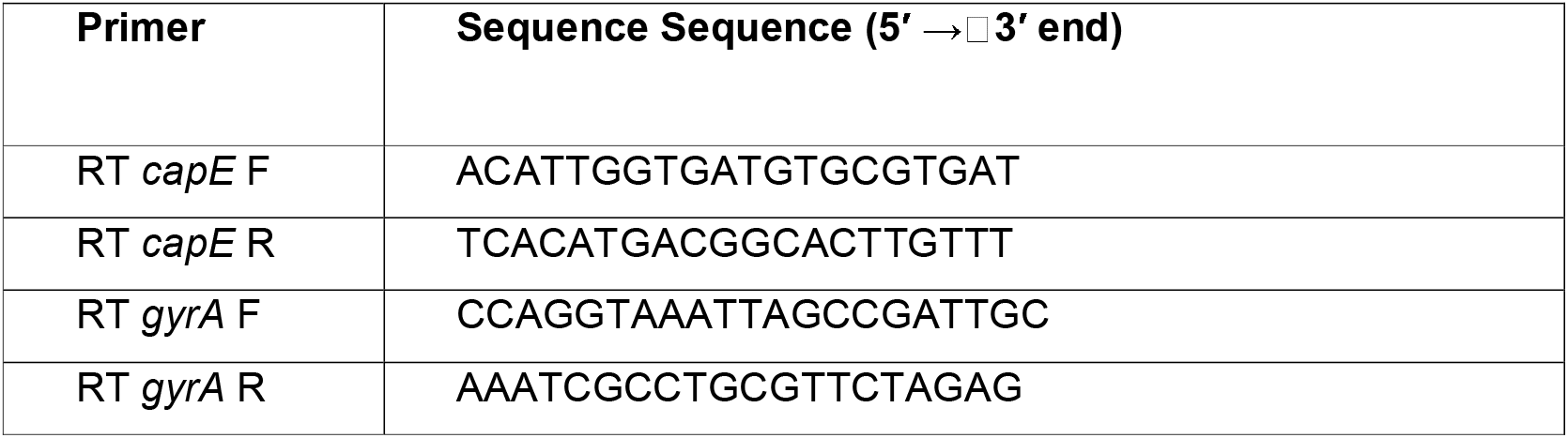
Oligonucleotide primers used in this study

**Table 2:** Clinical strains used in this study and the capsule production score.

| CC30 strain ID | Level of capsule production |  | CC22 strain ID | Level of capsule production |
| --- | --- | --- | --- | --- |
| ASARM88 | 1 |  | ASARM61 | 2 |
| ASARM90 | 1 |  | ASARM70 | 2 |
| ASARM92 | 1 |  | ASARM71 | 2 |
| ASARM63 | 0 |  | ASARM72 | 2 |
| ASARM112 | 0 |  | ASARM73 | 2 |
| ASARM147 | 2 |  | ASARM74 | 2 |
| ASARM156 | 0 |  | ASARM77 | 2 |
| ASARM161 | 2 |  | ASARM76 | 2 |
| ASARM180 | 2 |  | ASARM75 | 2 |
| ASARM185 | 2 |  | ASARM80 | 2 |
| ASARM190 | 2 |  | ASARM79 | 2 |
| ASARM197 | 2 |  | ASARM59 | 0 |
| ASARM210 | 2 |  | ASARM83 | 2 |
| ASARM213 | 1 |  | ASARM84 | 0 |
| ASASM4 | 2 |  | ASARM86 | 1 |
| ASASM22 | 1 |  | ASARM87 | 1 |
| ASASM35 | 2 |  | ASARM89 | 2 |
| ASASM6 | 2 |  | ASARM62 | 2 |
| ASASM43 | 2 |  | ASARM93 | 0 |
| ASASM46 | 2 |  | ASARM95 | 2 |
| ASASM47 | 2 |  | ASARM96 | 1 |
| ASASM8 | 2 |  | ASARM97 | 0 |
| ASASM59 | 0 |  | ASARM99 | 2 |
| ASASM14 | 0 |  | ASARM100 | 2 |
| ASASM17 | 2 |  | ASARM101 | 2 |
| ASASM66 | 2 |  | ASARM102 | 2 |
| ASASM67 | 2 |  | ASARM103 | 2 |
| ASASM78 | 2 |  | ASARM105 | 2 |
| ASASM79 | 2 |  | ASARM107 | 1 |
| ASASM89 | 2 |  | ASARM109 | 0 |
| ASASM98 | 1 |  | ASARM114 | 2 |
| ASASM99 | 2 |  | ASARM64 | 1 |
| ASASM100 | 2 |  | ASARM116 | 2 |
| ASASM103 | 0 |  | ASARM117 | 2 |
| ASASM116 | 2 |  | ASARM118 | 0 |
| ASASM131 | 1 |  | ASARM119 | 2 |
| ASASM133 | 2 |  | ASARM120 | 2 |
| ASASM134 | 1 |  | ASARM121 | 0 |
| ASASM147 | 1 |  | ASARM122 | 2 |
| ASASM149 | 2 |  | ASARM124 | 1 |
| ASASM156 | 0 |  | ASARM125 | 2 |
| ASASM157 | 0 |  | ASARM126 | 0 |
| ASASM159 | 2 |  | ASARM65 | 0 |
| ASASM166 | 1 |  | ASARM127 | 2 |
| ASASM126 | 2 |  | ASARM128 | 0 |
| ASASM173 | 2 |  | ASARM132 | 2 |
| ASASM174 | 2 |  | ASARM133 | 0 |
| ASASM200 | 0 |  | ASARM134 | 2 |
| ASASM201 | 2 |  | ASARM135 | 2 |
| ASASM202 | 2 |  | ASARM136 | 2 |
| ASASM212 | 2 |  | ASARM137 | 2 |
| ASASM220 | 2 |  | ASARM67 | 2 |
| ASASM237 | 1 |  | ASARM138 | 2 |
| ASASM250 | 2 |  | ASARM139 | 2 |
| ASASM268 | 2 |  | ASARM140 | 2 |
| ASASM272 | 1 |  | ASARM141 | 2 |
| ASASM281 | 2 |  | ASARM142 | 0 |
| ASASM285 | 2 |  | ASARM143 | 0 |
| ASASM288 | 2 |  | ASARM144 | 2 |
| ASASM289 | 0 |  | ASARM145 | 2 |
| ASASM292 | 1 |  | ASARM68 | 2 |
| ASASM293 | 2 |  | ASARM148 | 2 |
| ASASM297 | 2 |  | ASARM154 | 0 |
| ASASM303 | 2 |  | ASARM153 | 0 |
| ASASM307 | 2 |  | ASARM155 | 2 |
| ASASM320 | 1 |  | ASARM160 | 2 |
| ASASM329 | 2 |  | ASARM69 | 2 |
| ASASM330 | 1 |  | ASARM162 | 2 |
| ASASM331 | 2 |  | ASARM164 | 2 |
| ASASM340 | 0 |  | ASARM163 | 2 |
| ASASM345 | 2 |  | ASARM166 | 2 |
| ASASM351 | 0 |  | ASARM165 | 0 |
| ASASM358 | 0 |  | ASARM167 | 2 |
| ASASM369 | 0 |  | ASARM168 | 2 |
| ASASM391 | 1 |  | ASARM169 | 2 |
| ASASM395 | 2 |  | ASARM179 | 2 |
| ASASM406 | 2 |  | ASARM181 | 2 |
| ASASM407 | 2 |  | ASARM183 | 2 |
| ASASM408 | 2 |  | ASARM184 | 2 |
| ASARM436 | 0 |  | ASARM170 | 0 |
| ASASM443 | **2** |  | ASARM191 | **2** |
| ASASM445 | 2 |  | ASARM193 | 2 |
| ASASM446 | 0 |  | ASARM196 | 2 |
| ASASM451 | 1 |  | ASARM199 | 2 |
| ASASM457 | 2 |  | ASARM200 | 0 |
| ASASM458 | 1 |  | ASARM201 | 0 |
| ASASM373 | 2 |  | ASARM171 | 0 |
| ASASM377 | 2 |  | ASARM203 | 0 |
| ASASM378 | 2 |  | ASARM204 | 0 |
| ASASM379 | 2 |  | ASARM205 | 1 |
| ASASM385 | 2 |  | ASARM208 | 2 |
| EOE003 | 0 |  | ASARM209 | 2 |
| EOE023 | 0 |  | ASARM207 | 0 |
| EOE030 | 0 |  | ASARM211 | 2 |
| EOE035 | 0 |  | ASARM212 | 2 |
| EOE041 | 1 |  | ASARM172 | 2 |
| EOE042 | 0 |  | ASARMLT1 | 1 |
| EOE045 | 1 |  | ASARMLT2 | 1 |
| EOE052 | 1 |  | ASARMLT3 | 1 |
| EOE054 | 0 |  | ASARM195 | 2 |
| EOE057 | 1 |  | ASARM176 | 2 |
| EOE061 | 1 |  | ASARM177 | 2 |
| EOE065 | 1 |  | ASARM217 | 2 |
| EOE072 | 0 |  | ASARM220 | 2 |
| EOE073 | 1 |  | ASARM221 | 2 |
| EOE078 | 0 |  | ASARM222 | 0 |
| EOE083 | 1 |  | ASARM223 | 2 |
| EOE084 | 0 |  | ASARM224 | 2 |
| EOE086 | 1 |  | ASARM110 | 2 |
| EOE088 | 1 |  | ASARM108 | 2 |
| EOE089 | 1 |  | ASASM42 | 2 |
| EOE090 | 1 |  | ASASM56 | 2 |
| EOE091 | 1 |  | ASASM12 | 2 |
| EOE094 | 1 |  | ASASM61 | 2 |
| EOE096 | 0 |  | ASASM64 | 2 |
| EOE097 | 2 |  | ASASM71 | 2 |
| EOE098 | 2 |  | ASASM73 | 2 |
| EOE099 | 0 |  | ASASM90 | 0 |
| EOE100 | 1 |  | ASASM96 | 0 |
| EOE101 | 1 |  | ASASM97 | 2 |
| EOE102 | 0 |  | ASASM120 | 2 |
| EOE103 | 1 |  | ASASM132 | 2 |
| EOE104 | 0 |  | ASASM138 | 2 |
| EOE105 | 0 |  | ASASM140 | 2 |
| EOE106 | 1 |  | ASASM125 | 2 |
| EOE122 | 0 |  | ASASM127 | 2 |
| EOE125 | 1 |  | ASASM181 | 2 |
| EOE129 | 1 |  | ASASM186 | 2 |
| EOE130 | 1 |  | ASASM190 | 1 |
| EOE137 | 1 |  | ASASM246 | 0 |
| EOE140 | 1 |  | ASASM262 | 2 |
| EOE154 | 1 |  | ASASM390 | 2 |
| EOE155 | 1 |  | ASASM392 | 0 |
| EOE158 | 0 |  | ASASM398 | 2 |
| EOE118 | 0 |  | ASASM430 | 2 |
| EOE198 | 0 |  | ASASM168 | 2 |
| EOE205 | 1 |  |  |  |
| EOE220 | 1 |  |  |  |
| EOE120 | 0 |  |  |  |
| EOE225 | 1 |  |  |  |
| EOE233 | 1 |  |  |  |
| EOE234 | 1 |  |  |  |
| EOE237 | 1 |  |  |  |
| EOE268 | 2 |  |  |  |
| EOE269 | 2 |  |  |  |
| EOE274 | 1 |  |  |  |
| EOE275 | 1 |  |  |  |
| EOE276 | 1 |  |  |  |
| EOE029 | 0 |  |  |  |
| EOE161 | 1 |  |  |  |
| EOE162 | 0 |  |  |  |
| EOE163 | 2 |  |  |  |
| EOE165 | 2 |  |  |  |
| EOE166 | 2 |  |  |  |
| EOE167 | 1 |  |  |  |
| EOE169 | 1 |  |  |  |
| EOE171 | 1 |  |  |  |
| EOE173 | 0 |  |  |  |
| EOE174 | 0 |  |  |  |
| EOE175 | 1 |  |  |  |
| EOE176 | 0 |  |  |  |
| EOE208 | 0 |  |  |  |
| EOE126 | 0 |  |  |  |
| EOE229 | 1 |  |  |  |

### Selection and verification of SCV strains

*S. aureus* strain Newman was grown in TSB at 37°C incubated in shaking incubator overnight. The culture was diluted 1/10 into TSB with 2 μg/ml gentamicin and incubated for 8 h. The resulting culture was then plated on blood agar containing 2 μg/ml gentamicin. Pin-prick sized colonies were further isolated by streaking onto fresh agar plates with 2 μg/ml gentamicin. Auxotrophy to both menadione and hemin was examined by placing a filter disk saturated in these growth reagents onto a freshly inoculated lawn of the purified SCV colonies, and enhanced growth surrounding the disk visually examined.

## Results and Discussion

### Capsule production varies across closely related *S. aureus* bacteraemia isolates

Recent work has suggested that there is significant variability amongst clinical *S. aureus* isolates in the amount of capsule they produce [6, 7]. Given the importance of capsule in protecting the bacteria from many aspects of the human immune system we sought to examine the variability of this in isolates from invasive disease, where the anti-bacterial effects of the immune system should be the most stringent. We focussed on a collection of isolates from 300 cases of bacteraemia, representing both the two major clones of MRSA strains circulating in UK and Irish hospitals (CC22 and CC30), as well as the two major capsule serotype of *S. aureus* that cause disease in humans (capsule type 5 (CP5) and type 8 (CP8)). We performed a semi-quantification of capsule production by each isolate using anti-CP5 (for the CC22 isolates) and anti-CP8 (for the CC30 isolates) antiserum. The reactivity of the antisera was demonstrated using a pair of wild type and capsule negative isogenic mutants (Fig. 1). Across the clinical bacteraemia isolates there was significant variability in capsule production, with no detectable capsule being produced by 23% of the isolates (fig. 1, Table 1).

**Figure 1:**
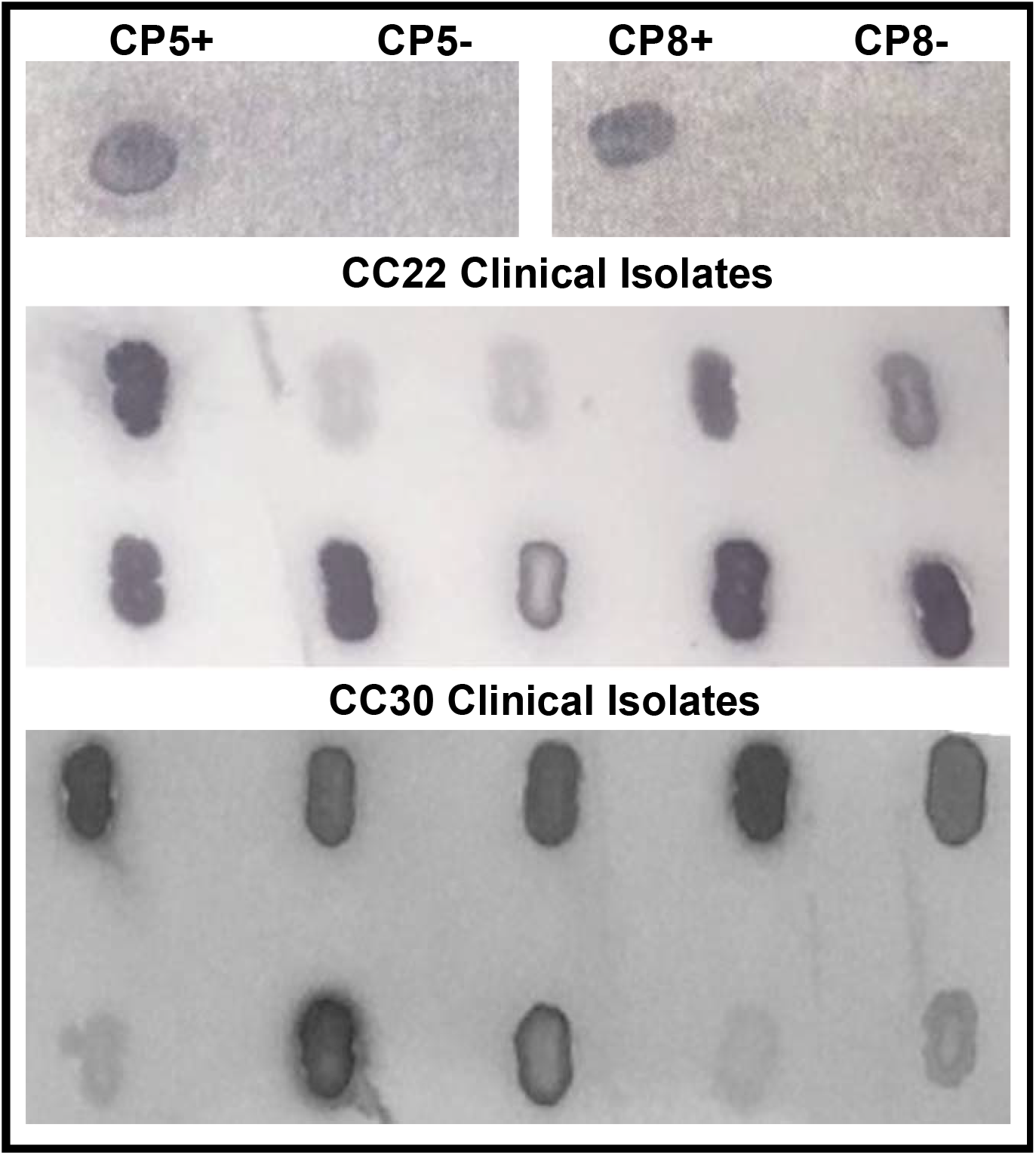
Capsule production varies significantly across clinical bacteraemia isolates. Immunoblots of *S. aureus* isolates were performed with either anti-CP5 or anti-CP8 antiserum. Blots of isogenic CP5 and CP8 wild type and cap-mutant were performed as controls (top row). Ten CC22 (rows 2 & 3) and 10 CC30 (rows 4 &5) isolates representative of the variability in intensity of anti-capsule anti-serum binding are presented. The CC22s were probed with the anti-CP5 antiserum and the CC30s with the anti-CP8 antiserum.

### The genetic basis of the variability in capsule production is multifactorial

As the genome sequence for each of the 300 clinical *S. aureus* isolates was available we performed a GWAS (genome wide association study) to identify polymorphic loci that associated with the level of capsule produced by the isolates. For this, the data from the two distinct clones were analysed independently, with population structure within the clones being accounted for (Fig. 2, Tables 3 & 4). We applied both uncorrected and corrected (for multiple comparisons) significance thresholds to this analysis, as our previous work has demonstrated that the stringency of multiple correction approaches increases the likelihood of type II errors, or false negative results. Only one locus was found associated at the multiple test corrected significance threshold: the *agrC* gene, which is part of a well-established regulatory system of many virulence factors including capsule [9], providing good proof of concept for this approach. A further 169 loci were found associated with capsule production at the P<0.05 significance threshold, including two genes, in which mutations result in the switching of *S. aureus* to the small colony variant (SCV), or persistor phenotype: *fabD* and *menD*. SCVs auxotrophic for fatty acids that are more resistant to FAS-II inhibitors, such as triclosan, are associated with mutations in the *fabD* gene [22]; whereas SCVs auxotrophic for menadione that are more resistant to aminoglycoside antibiotics, such as gentamicin, are associated with mutations in the *menD* gene [15, 16].

**Table 3:** Loci associated with capsule production in the CC22 collection of *S. aureus* isolates.

| Gene or Locus Tag | Protein function | P value |
| --- | --- | --- |
| SAEMRSA15_RS00260 | type I restriction endonuclease subunit R | 0.00014021 |
| Intergenic between<br>SAEMRSA15_RS13970 and *clfB* |  | 0.00034659 |
| SAEMRSA15_RS11275 | thiol-disulfide oxidoreductase DCC family protein | 0.00034659 |
| SAEMRSA15_RS12900 | NAD(P)-dependent oxidoreductase | 0.00034659 |
| SAEMRSA15_RS01030 | type 1 glutamine amidotransferase | 0.00034659 |
| SAEMRSA15_RS00265 | hypothetical protein | 0.00034659 |
| SAEMRSA15_RS08245 | acetyl-CoA carboxylase biotin carboxylase subunit | 0.00034659 |
| SAEMRSA15_RS10695 | LytTR family DNA-binding domain-containing protein | 0.00075253 |
| SAEMRSA15_RS10690 | GHKL domain-containing protein | 0.00077225 |
| SAEMRSA15_RS02555 | RNA polymerase sigma factor | 0.00120048 |
| *ileS* | isoleucine--tRNA ligase | 0.00139006 |
| SAEMRSA15_RS12840 | glycerate kinase | 0.00280998 |
| SAEMRSA15_RS11120 | ATP synthase subunit I | 0.0030996 |
| SAEMRSA15_RS02630 | amidohydrolase | 0.00366703 |
| SAEMRSA15_RS12955 | APC family permease | 0.00386528 |
| SAEMRSA15_RS13455 | LrgB family protein | 0.00400753 |
| *menD* | 2-succinyl-5-enolpyruvyl-6-hydroxy-3-cyclohexene-1-carboxylic-acid synthase | 0.00507518 |
| SAEMRSA15_RS09800 | metal-dependent hydrolase | 0.00516556 |
| *graR* | response regulator transcription factor<br>GraR/ApsR | 0.00546077 |
| *rsmG* | 16S rRNA (guanine(527)-N(7))-methyltransferase RsmG | 0.00895798 |
| Intergenic between<br>SAEMRSA15_RS08855 and<br>SAEMRSA15_RS08860 |  | 0.01032749 |
| Intergenic between<br>SAEMRSA15_RS01390 and *brnQ* |  | 0.01032749 |
| *sbnC* | staphyloferrin B biosynthesis protein SbnC | 0.01032749 |
| SAEMRSA15_RS03060 | WecB/TagA/CpsF family<br>glycosyltransferase | 0.0108 |
| Intergenic between *guaC* and<br>SAEMRSA15_RS06385 |  | 0.01141554 |
| SAEMRSA15_RS04275 | Glu/Leu/Phe/Val dehydrogenase | 0.01534142 |
| SAEMRSA15_RS14050 | amidase domain-containing protein | 0.01534142 |
| SAEMRSA15_RS04670 | ATP-binding cassette domain-containing<br>protein | 0.01534142 |
| SAEMRSA15_RS08370 | histidine--tRNA ligase | 0.01534142 |
| SAEMRSA15_RS08350 | replication-associated recombination<br>protein A | 0.01534142 |
| Intergenic between<br>SAEMRSA15_RS02335 and *tilS* |  | 0.01534142 |
| SAEMRSA15_RS14225 | ATP phosphoribosyltransferase | 0.01534142 |
| SAEMRSA15_RS11600 | hypothetical protein | 0.01534142 |
| SAEMRSA15_RS09105 | MFS transporter | 0.01577408 |
| SAEMRSA15_RS11705 | energy-coupling factor transporter ATPase | 0.01652847 |
| SAEMRSA15_RS13060 | hypothetical protein | 0.01671052 |
| *sdhA* | succinate dehydrogenase flavoprotein subunit | 0.01836527 |
| *fabD* | ACP S-malonyltransferase | 0.01905729 |
| *ribB* | 3,4-dihydroxy-2-butanone-4-phosphate synthase | 0.02115675 |
| SAEMRSA15_RS13160 | DUF3427 domain-containing protein | 0.02147993 |
| SAEMRSA15_RS02320 | nucleotide pyrophosphohydrolase | 0.02182662 |
| Intergenic between brnQ and SAEMRSA15_RS00800 |  | 0.02388376 |
| *radA* | DNA repair protein RadA | 0.02388376 |
| SAEMRSA15_RS02405 | tRNA-Lys | 0.02388376 |
| SAEMRSA15_RS01035 | PrsW family intramembrane metalloprotease | 0.0245801 |
| SAEMRSA15_RS01275 | ABC transporter permease | 0.02728306 |
| SAEMRSA15_RS07140 | hypothetical protein | 0.02954242 |
| *pyk* | pyruvate kinase | 0.0303263 |
| SAEMRSA15_RS11310 | hypothetical protein | 0.03044146 |
| *feoB* | ferrous iron transport protein B | 0.03071363 |
| SAEMRSA15_RS02010 | YbcC family protein | 0.03520898 |
| SAEMRSA15_RS01135 | CDP-glycerol glycerophosphotransferase family protein | 0.03607997 |
| *dltB* | PG:teichoic acid D-alanyltransferase DltB | 0.03944867 |
| SAEMRSA15_RS05405 | YfcC family protein | 0.04425032 |
| SAEMRSA15_RS05040 | DUF4064 domain-containing protein | 0.04807582 |
| SAEMRSA15_RS03910 | thermonuclease family protein | 0.04901599 |

**Table 4:** Loci associated with capsule production in the CC30 collection of *S. aureus* isolates.

| Gene or Locus Tag | Protein function | P value |
| --- | --- | --- |
| *agrC* | autoinducer sensor protein | 4.06 x 10<sup>-5</sup> |
| SAR1756 | hypothetical protein | 0.000610781 |
| *kdpA* | putative potassium-transporting ATPase a chain | 0.00079868 |
| *leuB* | 3-isopropylmalate dehydrogenase | 0.00079868 |
| SAR2555 | conserved hypothetical protein | 0.001102934 |
| Intergenic between *sstD* and SAR0791 |  | 0.002038995 |
| *ccpA* | catabolite control protein A | 0.005730767 |
| SAR2382 | putative transcriptional regulator | 0.005772766 |
| SAR2759 | putative aminotransferase-putative imidazoleglycerol-phosphate dehydratase | 0.005813137 |
| SAR1218 | putative membrane protein | 0.007268775 |
| SAR0457a | hypothetical protein | 0.008232546 |
| SAR2533 | putative ketopantoate reductase | 0.008799502 |
| SAR0109 | putative transporter protein | 0.009570567 |
| *yycG* | Two-component regulatory system family, sensor kinase protein. | 0.010100134 |
| *thiE* | putative thiamine-phosphate pyrophosphorylase | 0.010702445 |
| *fabD* | putative malonyl CoA-acyl carrier protein transacylase | 0.011824195 |
| SAR1343 | amino acid permease | 0.01386841 |
| SAR2522 | putative glycerate kinase | 0.01579356 |
| SAR1674 | putative GTPase | 0.018539766 |
| SAR0122 | putative transport protein | 0.018645214 |
| SAR1668 | conserved hypothetical protein | 0.019293181 |
| *ilvA* | threonine dehydratase biosynthetic | 0.020495129 |
| *dfrB* | dihydrofolate reductase type I | 0.021075917 |
| Intergenic between *polS* and *proC* |  | 0.022922752 |
| SAR2025 | putative ABC transporter ATP-binding protein | 0.02367155 |
| SAR0793 | hypothetical protein | 0.023724266 |
| SAR1619 | putative exported protein | 0.023978842 |
| SAR0743 | putative sodium:sulfate symporter protein | 0.02413635 |
| *arsR2* | arsenical resistance operon repressor 2 | 0.024274209 |
| SAR0108 | putative peptidase | 0.024560717 |
| SAR0559 | putative aminotransferase | 0.02466874 |
| SAR1002 | putative membrane protein | 0.02622901 |
| SAR0942 | putative membrane protein | 0.026651981 |
| SAR2740 | conserved hypothetical protein | 0.026988447 |
| *ureC* | urease alpha subunit | 0.027434322 |
| *qoxB* | putative quinol oxidase polypeptide I | 0.028188487 |
| *mnhD* | Na<sup>+</sup>/H<sup>+</sup> antiporter subunit | 0.030064463 |
| SAR2534 | putative transport protein | 0.030662069 |
| SAR2779 | putative N-acetyltransferase | 0.031463295 |
| SAR1684 | conserved hypothetical protein | 0.031847864 |
| SAR1699 | conserved hypothetical protein | 0.031847864 |
| SAR1995 | putative lipoprotein | 0.031847864 |
| SAR0463 | putative lipoprotein | 0.031847864 |
| SAR0010 | putative membrane protein | 0.033712843 |
| SAR0245 | putative zinc-binding dehydrogenase | 0.035487178 |
| *ureE* | urease accessory protein UreE | 0.035624064 |
| *odhA* | 2-oxoglutarate dehydrogenase E1 component | 0.036566931 |
| SAR2186 | conserved hypothetical protein | 0.037233673 |
| *hisB* | putative imidazoleglycerol-phosphate dehydratase | 0.037571061 |
| SAR0291 | putative membrane protein | 0.038246027 |
| SAR1876 | hypothetical protein | 0.038551026 |
| SAR1703 | putative oxygenase | 0.039218922 |
| SAR1655 | putative methyltransferase | 0.039424074 |
| SAR2464 | TetR family regulatory protein | 0.039515902 |
| SAR0987 | conserved hypothetical protein | 0.039515902 |
| SAR1868 | aldo/keto reductase family protein | 0.039515902 |
| SAR0466 | MutT domain containing protein | 0.039515902 |
| SAR0278 | putative exported protein | 0.039515902 |
| Intergenic between *rsbU* and SAR2156 |  | 0.039515902 |
| *ldh1* | L-lactate dehydrogenase 1 | 0.039515902 |
| *mvaD* | mevalonate diphosphate decarboxylase | 0.039515902 |
| SAR0969 | conserved hypothetical protein | 0.039515902 |
| SAR1973 | putative membrane protein | 0.039515902 |
| SAR0247 | putative zinc-binding dehydrogenase | 0.039515902 |
| SAR0770 | conserved hypothetical protein | 0.039515902 |
| SAR2619 | thiamine pyrophosphate enzyme | 0.039515902 |
| SAR1281 | conserved hypothetical protein | 0.039515902 |
| SAR0655 | putative Na<sup>+</sup> dependent nucleoside transporter | 0.039515902 |
| SAR1332 | response regulator | 0.039515902 |
| SAR2588 | putative membrane protein | 0.039515902 |
| SAR1165 | hypothetical protein | 0.039515902 |
| SAR1221 | putative CoA synthetase protein | 0.039515902 |
| Intergenic between SAR0994 and tRNA-Ser |  | 0.039515902 |
| SAR2006 | conserved hypothetical protein | 0.039515902 |
| SAR0636 | putative membrane protein (pseudogene) | 0.039515902 |
| SAR2780 | putative membrane protein | 0.039515902 |
| SAR1141 | Similar to Staphylococcus aureus exotoxin | 0.039515902 |
| SAR1670 | conserved hypothetical protein | 0.039515902 |
| SAR1685 | putative biotin carboxylase subunit of acetyl-CoA carboxylase | 0.039515902 |
| *lysS* | lysyl-tRNA synthetase | 0.039515902 |
| Intergenic between *lysS* and SAR1413 |  | 0.039515902 |
| Intergenic between SAR1326 and SAR1327 |  | 0.039515902 |
| *pbp4* | penicillin-binding protein 4 | 0.039515902 |
| SAR0880 | conserved hypothetical protein | 0.039515902 |
| Intergenic between *ehb* and SAR1448 |  | 0.039515902 |
| *vraD* | ABC transporter ATP-binding protein | 0.039515902 |
| SAR1265 | putative pyruvate<br>flavodoxin/ferredoxin oxidoreductase | 0.039515902 |
| SAR0147 | putative nucleotidase | 0.039515902 |
| Intergenic between *sodM*<br>and *sasG* |  | 0.039515902 |
| SAR1193 | hypothetical protein | 0.039515902 |
| SAR0509 | putative RNA binding protein | 0.039515902 |
| Intergenic between<br>SAR1932 and SAR1933 |  | 0.039515902 |
| SAR0810 | putative phosphohydrolase | 0.039515902 |
| Intergenic between *rpmH*<br>and *dnaA* |  | 0.039515902 |
| *bglA* | 6-phospho-beta-glucosidase | 0.039515902 |
| Intergenic between<br>SAR1870 and a SAM<br>riboswitch |  | 0.039515902 |
| *cysJ* | putative sulfite reductase [NADPH]<br>flavoprotein alpha-component | 0.039515902 |
| SAR2424 | putative aldose 1-epimerase | 0.039515902 |
| *pta* | putative phosphate acetyltransferase | 0.039515902 |
| SAR0269a | hypothetical protein | 0.039515902 |
| tRNA-Thr | tRNA Thr anticodon TGT, Cove score<br>85.17 | 0.039515902 |
| SAR1352 | putative transketolase | 0.039515902 |
| SAR0290 | hypothetical protein | 0.039515902 |
| *crtN* | squalene synthase | 0.039515902 |
| SAR1840 | putative exported protein | 0.039630048 |
| SAR1941 | RNA pseudouridylate synthase | 0.041094881 |
| SAR2411 | putative transport protein | 0.041580291 |
| SAR0900 | putative pyridine nucleotide-<br>disulphide oxidoreductase | 0.042472995 |
| *folC* | putative folylpolyglutamate synthase | 0.042664521 |
| *sasC* | putative surface anchored protein | 0.043300254 |
| SAR2119 | membrane anchored protein | 0.044028342 |
| SAR1279 | conserved hypothetical protein | 0.047982904 |
| *gidB* | putative glucose inhibited division<br>protein B | 0.048056799 |
| SAR2787 | hypothetical protein | 0.048327331 |

**Figure 2:**
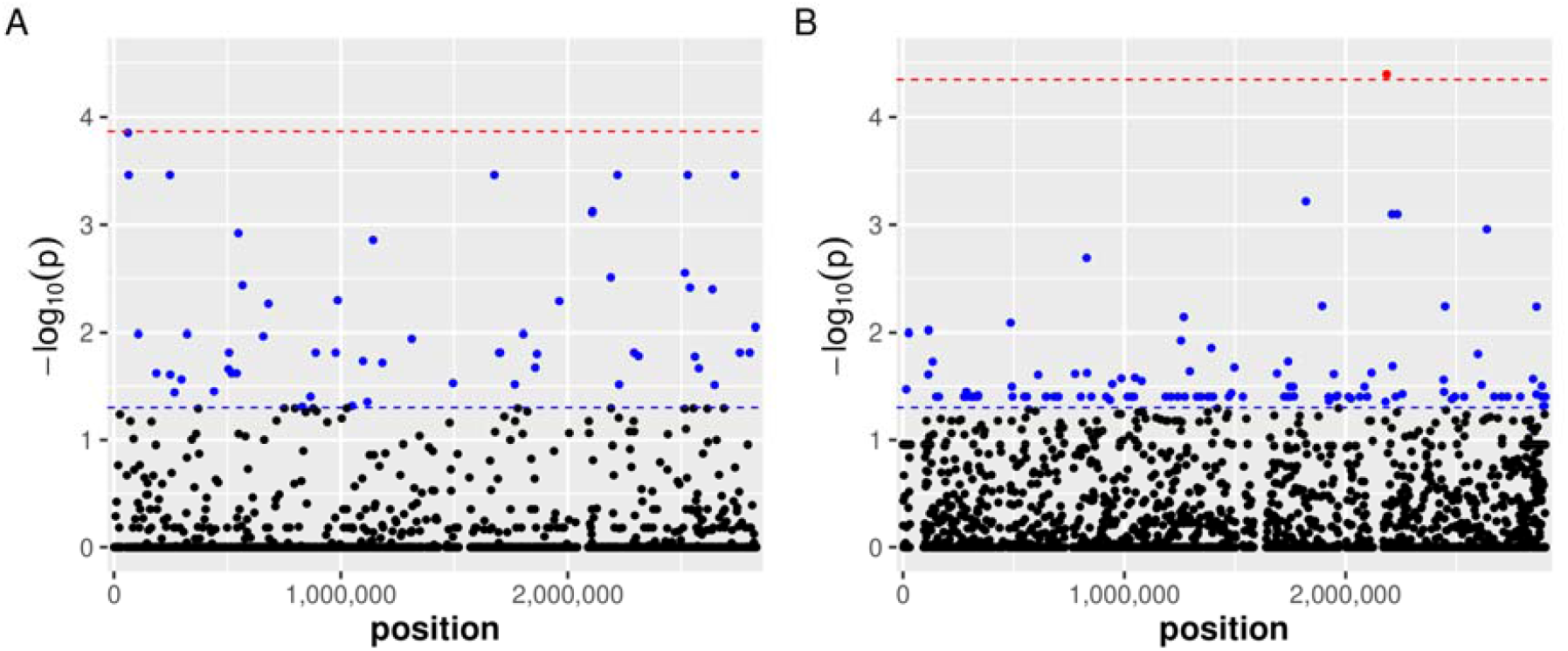
*S. aureus* loci associated with capsules production. Manhattan plots representing the results of a GWAS analysis identifying polymorphic loci associated with the level of capsule produced by (A) 136 CC22 and (B) 159 CC30 bacteraemia isolates. The x-axis represent the genomic position of the polymorphisms relative to the origin of replication and the y axis represents the strength of the association with capsule production. Uncorrected (P<0.05) and multiple tests corrected (P<1.3×10^−4^, for CC22; and P<4.5×10^−5^ for CC30s) significance thresholds are indicated as blue and red lines, respectively.

### Functional verification of the role of *menD* in capsule production

There are contradictory reports in the literature on the effect the switch to SCV has on capsule production [19-21], and as such we sought to resolve these contradictions by verifying our GWAS findings with a focus on the *menD* gene. The *menD* gene encodes an enzyme involved in the biosynthesis of menadione, which is a vitamin K2 precursor that is synthesised by *S. aureus* [15]. The importance of menadione for efficient respiration by the bacteria is such that inactivation of the gene results in a slow-growing small colony variant (SCV) phenotype [15, 16]. There are other metabolic pathways that can mutate and result in an SCV phenotype such as in the hemin biosynthesis pathway [18], and collectively the SCV phenotype is associated with significant changes in *S. aureus* virulence, in particular with regards to reduced toxin production [18].

Given our association between polymorphisms in the *menD* gene and capsule production in two distinct clones of *S. aureus*, we sought to examine this in further detail. SCVs were selected from a culture *S. aureus* strain Newman by overnight growth in gentamicin (2μg/ml), on the basis of their enhanced resistance to the aminoglycoside class of antibiotics. Of these SCVs we identified a menadione auxotrophic SCV, as well as a hemin auxotrophic SCV as a comparator by restoring the growth defect through the addition of either menadione or hemin on a disc (Fig. 3a). We performed immunoblots of the wild type strain Newman and the SCVs, where there was a significant effect on capsule production for the menadione auxotrophic SCV, but not the hemin auxotrophic SCV (Fig. 3b and c). To further examine the effect on capsule production, we quantified the transcription of the *capE* gene, where we found this to be significantly reduced in the menadione auxotroph, but not the hemin auxotroph (Fig. 3d). While further work is underway to examine the effect mutations in *fabD* and triclosan resistance has on capsule production, here we have verified the observed association between the *menD* gene and capsule production. The discrepancy between the levels of capsule production by the *hemB* and *menD* SCVs may also explain some of the discrepancy in the literature in relation to capsule production by SCVs, in that the effect is dependent upon the pathway that becomes mutated.

**Figure 3:**
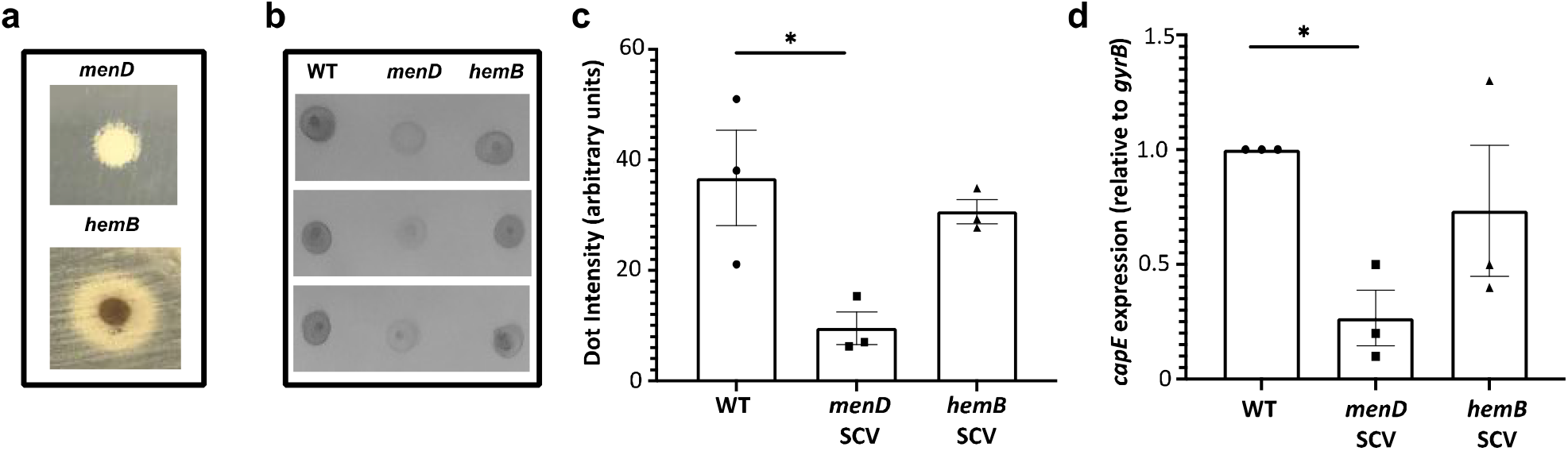
Capsule production is affected in a menadione auxotrophic SCV. (**a**) *menD* and *hemB* SCVs of *S. aureus* strain Newman were selected, and auxotrophy to menadione and hemin determined by examining enhanced growth of the SCV when the medium was supplemented with a disk containing the respective growth reagent. (**b & c**) Immunoblotting of the wild type Newman and the *menD* and *hemB* SCVs demonstrate that the capsule production is only affected in the menadione auxotrophic SCV. (**d**) Transcription of the *capE* gene is lower in the menadione-auxotrophic SCV relative wild type Newman, but not in the hemin auxotroph.

### The *menD* polymorphisms in the clinical isolates do not affect growth but do increase resistance to gentamicin

Having demonstrated that capsule production is affected in the menadione auxotrophic SCVs, we examined whether the isolates with polymorphisms within our collections of bacteraemia isolates also had the SCV phenotype. There were nine isolates with non-synonomous polymorphism in the *menD* gene, and the position and effect of the SNPs on the amino acids sequence are illustrated in Fig. 4. We selected at random nine isolates from the collection with the non-polymorphic *menD* gene (i.e. identical to the respective reference strains MRSA252 [23] and HO 5096 0412 [24]). These isolates were grown in TSB with and without 2μg/ml of gentamicin to examine the two main features of SCVs, slow growth and increased resistance to gentamicin. We found that the clinical *menD* variants grew as well as those with the reference *menD* gene in TSB, demonstrating that they have no growth defect *in vitro*. However, in the presence of gentamicin we found that the variants had a growth advantage, which suggests they have a partial SCV phenotype, at least with respect to their enhanced resistance to this antibiotic (fig. 5).

**Figure 4:**
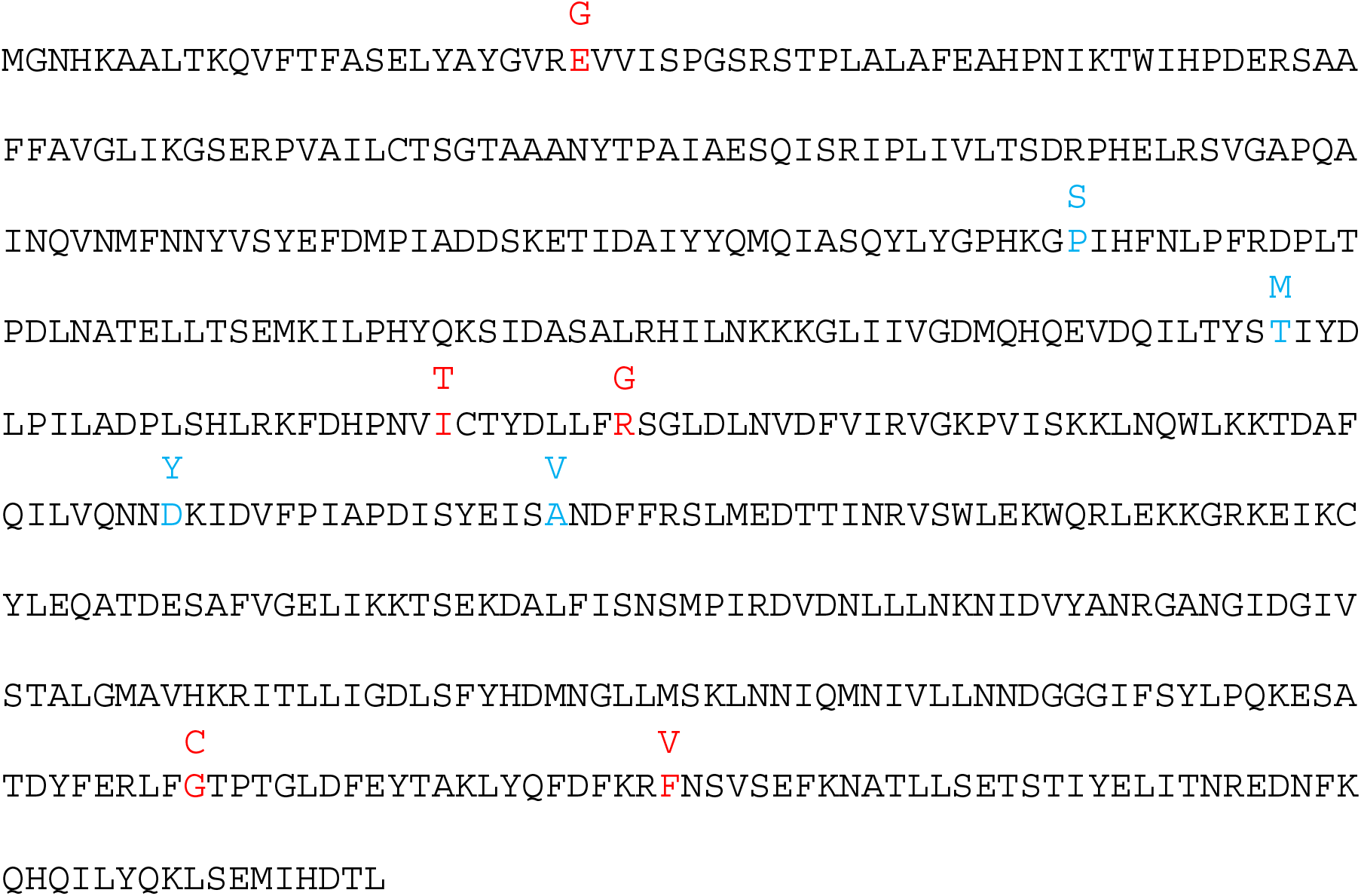
The *S. aureus* MenD amino acid sequence. The effect of the non-synonomous polymorphism present in the CC30 (indicated in blue font) and CC22 (in red font) collection of isolates studied here are indicated.

**Figure 5:**
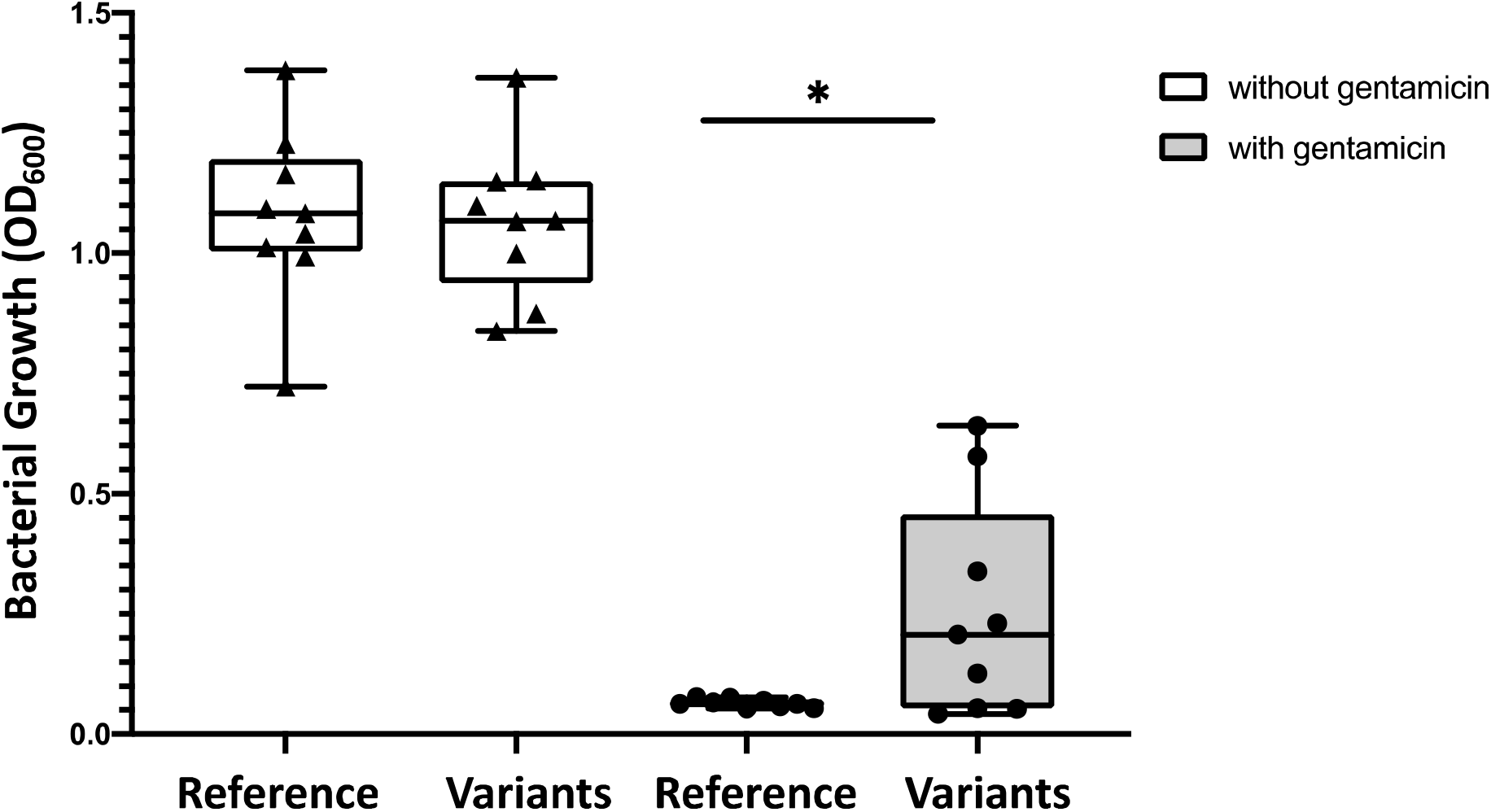
Clinical isolates with *menD* polymorphisms have no growth defect but are more resistant to gentamicin. The growth of the nine clinical isolates containing non-synonomous polymorphism in the *menD* gene was compared to that of nine randomly selected isolates with the wild type or reference *menD* gene. In TSB we observed no effect on growth associated with the polymorphic *menD* gene, however in a concentration of 2μg/ml of gentamicin, the *menD* variants grew significantly better.

In summary, in this study we have identified novel putative effectors of capsule production by *S. aureus*, including the menadione biosynthesis pathway. In doing so we have resolved an apparent contradiction in the literature with respect to the effect that the switch from normal growth to the SCV form has on capsule production. We found that this crucially depends on which metabolic pathway has been mutated to result in the switch. What is intriguing is that all isolates studied here were from cases of bacteraemia, and despite the importance of capsule production to the protection of the bacteria from many aspects of the human immune system, we found that around 1 in 5 isolates do not express capsule to any detectable levels. It is possible that the loss of capsule coincides with enhanced antibiotic resistance, as we have observed here for mutations in *menD*. With further investigation we may find that mutations of the other associated loci also confer advantages to the bacteria that over-ride the cost associated with the loss of capsule. But what is clear is that even within a clone, *S. aureus* is a highly adaptable and diverse in its means of causing disease, which may explain our lack of success in producing an effective vaccine using capsule as its major target.

## Authors Statements

### Contributions

DA developed the methodology, performed experiments, analyzed data and contributed to writing the manuscript. TB provided supervisory support, analyzed data and contributed to writing the manuscript. AE & JL provided resources and contributed to writing the manuscript. MR analyzed data and contributed to writing the manuscript. RCM conceptualized the projects, provided supervisory support, analyzed data and contributed to writing the manuscript.

### Conflict of Interests

we the authors declare that we have no conflicts of interest associated with the work described in this manuscript.

### Funding information

this work was funded by a PhD studentship to DA funded by the Saudi Arabian Cultural Bureau and RCM is a Wellcome Trust funded Investigator (Grant reference number: 212258/Z/18/Z)

## Notes

### Competing Interest Statement

The authors have declared no competing interest.

